# Same-sex mounting in single-housed C57BL/6J female mice is not coupled to estrous state

**DOI:** 10.1101/2025.07.01.662591

**Authors:** Cassidy A. Malone, Xin Zhao, Selina B. Xu, Katherine A. Tschida

## Abstract

Female-female mounting is widespread in mammals. Although the factors that regulate the display of female-female mounting likely vary according to species and to behavioral context, a large body of work has investigated the relationship of female-female mounting to female hormonal status. While this relationship has been well explored in ovariectomized and hormonally manipulated female rodents, less is known about the potential relationship between female-female mounting and estrous state in naturally cycling female rodents. Recent studies found that short-term social isolation robustly promotes same-sex mounting in C57BL/6J female mice. In the current study, we tested whether displays of female-female mounting by naturally cycling single-housed C57BL/6J females during interactions with naturally cycling, group-housed stimulus females depend on the estrous state of either female (n = 34 independent pairs). We found that the likelihood of mounting by single-housed females did not differ according to their estrous state. Moreover, single-housed females displayed mounting in trials that included receptive stimulus females, as well as in trials that included non-receptive stimulus females, and same-sex mounting was observed across all resident-stimulus female estrous state combinations. These findings indicate that same-sex mounting by single-housed C57BL/6J females is not coupled to the estrous state of either female in the pair. Future work remains to determine the factors that regulate the occurrence of female-female mounting in this species and behavioral context.

## Introduction

Female-female mounting is widespread throughout the animal kingdom and has been observed in mammals ranging from primates to rodents (Bailey and Zuk, 2009; Dagg, 1984; Gómez et al., 2023). The factors that regulate female-female mounting likely vary according to species, as well as behavioral and developmental context. Nonetheless, given the role of male-female mounting in mammalian courtship behavior, a large body of work has considered the possibility that female-female mounting reflects some aspect of female sexual behavior, and correspondingly, has explored the relationship of female-female mounting to female hormonal status. These past works have been conducted most extensively in rodents, including rats (Baum et al., 1974; Beach and Rasquin, 1942; Beach, 1968; Coniglio and Clemens, 1972; de Jonge et al., 1986; Fang and Clemens, 1999; Södersten, 1972; Van de Poll et al., 1988), hamsters (Noble, 1974, 1977), and mice (Bakker et al., 2006, 2002; Edwards and Burge, 1971; Martel and Baum, 2009; Wersinger et al., 1997; Williamson et al., 2019).

The literature regarding the relationship in female rodents between hormonal status and the display of same-sex mounting has been dominated by studies using ovariectomized (OVX) and hormonally manipulated females, with relatively fewer studies using naturally cycling females. Early work in rats reported that OVX females engaged in same-sex mounting at rates similar to gonadally intact controls (Beach and Rasquin, 1942; Beach, 1968). However, it was also reported that OVX female rats displayed lower rates of same-sex mounting than gonadally intact females, and that subsequent estradiol administration promoted female-female mounting (Södersten, 1972). Additional studies reported that OVX female rats and hamsters treated with estradiol or testosterone display elevated rates of same-sex mounting relative to OVX controls (Baum et al., 1974; de Jonge et al., 1986; Noble, 1974, 1977). Similarly, OVX female mice treated with testosterone display elevated rates of same-sex mounting (Bakker et al., 2002, 2002; Edwards and Burge, 1971; Martel and Baum, 2009; Wersinger et al., 1997), and these effects were attenuated in estrogen receptor alpha knock-out females (Wersinger et al., 1997), in aromatase knock-out females (Bakker et al., 2002), as well as in females lacking the gene for alpha-fetoprotein, an estrogen-binding fetal plasma protein (Bakker et al., 2006).

As for the relationship of mounting to the hormonal status of a stimulus female, past studies have reported that female rats are more likely to mount sexually receptive stimulus females (i.e., OVX females hormonally induced to be sexually receptive) than non-receptive (OVX) stimulus females (Coniglio and Clemens, 1972; Fang and Clemens, 1999; Van de Poll et al., 1988). Subsequently, many studies of same-sex mounting in female rodents include hormonally-induced stimulus females by default (Bakker et al., 2006, 2002; Baum et al., 1974; de Jonge et al., 1987; de Jonge et al., 1986; Edwards and Burge, 1971; Martel and Baum, 2009; Noble, 1974, 1977; Södersten, 1972; Wersinger et al., 1997). Together, these studies suggest that hormonal status can influence same-sex mounting in OVX female rodents, but the relationship of estrous state to same-sex mounting in naturally cycling female rodents is less clear.

In mice, there are relatively few reports of same-sex mounting in naturally cycling females. However, recent studies found that housing status strongly regulates the occurrence of same-sex mounting in female mice. A study using C57BL/6N mice found that the majority of females (6 of 10) that had been single-housed for at least 7 days mounted a socially novel stimulus female (Hashikawa et al., 2017). Likewise, work in C57BL/6J mice found that although female-female mounting was not observed in pairs including two group-housed females, the majority of single-housed females displayed same-sex mounting during an interaction with a novel stimulus female (Zhao et al., 2021; 2025). In all three studies, most (≥ 50%) but not all single-housed females displayed mounting, and the factors that regulate whether a given single-housed female displays mounting remain unknown. Based on the literature implicating hormonal status as a factor that can regulate same-sex mounting in female rodents, we conducted the current study to test whether displays of same-sex mounting in single-housed C57BL/6J females are coupled to the estrous state of either the single-housed female or the stimulus female.

## Materials and Methods

### Ethical Note

All experiments and procedures were conducted according to protocols approved by the Cornell University Animal Care and Use Committee (protocol #2020-001).

### Subjects

Adult (n = 68, > 8 weeks old) female C57BL/6J (Jackson Laboratories, 000664) mice were group-housed with 2–3 female siblings from weaning until the start of experiments. Mice were kept on a 12h:12h reversed light/dark cycle and given ad libitum food and water for the duration of the experiment. A running wheel (Innovive) was present in all home cages from weaning onward and was removed immediately before the social interaction test. Female subjects did not have social interactions with novel mice (female or male) prior to the start of experiments.

### Study design

Female subject mice were either individually housed in clean cages for three days prior to behavioral testing (single-housed resident females) or group-housed with same-sex siblings prior to testing (group-housed stimulus females). On the day of the social interaction test, a single-housed resident was placed in her home cage within a sound-attenuating recording chamber (Med Associates). A custom lid (with tall sides and no top) was placed on the cage to allow video and audio recordings. The recording chamber was equipped with an infrared light source (Tendelux) and a webcam (Logitech, with the infrared filter removed to enable video recording under infrared lighting). A socially novel (non-littermate) group-housed stimulus female was then placed in the single-housed resident’s home cage, and the pair was allowed to interact freely for 20 minutes. Each female was only used in a single trial, and a total of n = 68 females were used in the study (n = 34 as single-housed residents and n = 34 as group-housed stimulus females).

### Measurements of female estrous state

Vaginal cytology was evaluated for single-housed resident females and group-housed stimulus females on both the day before and the day of the social interaction test, to ensure accurate estrous staging. On the day of the social interaction test, vaginal cytology was collected for both female subjects immediately following the social interaction test. Vaginal cytology samples were collected by vaginal lavage with PBS (50 μL). Samples were pipetted directly onto slides and allowed to dry. After drying, slides were stained with Jorvet Dip Quick Stain (Jorgensen Laboratories) according to the manufacturer instructions. Stained slides were imaged at 20x magnification on a light microscope, and estrous state was determined by cell proportions as outlined previously (Cora et al., 2015).

### Scoring of behaviors from video recordings

Trained observers blinded to female estrous state results used BORIS v.8.13 to score times in webcam videos during which the single-housed resident female mounted the group-housed stimulus female. We did not observe any instances of group-housed stimulus females mounting single-housed residents.

### Statistical analyses

Fisher’s exact tests were used to compare the proportions of trials with mounting (alpha = 0.05). No statistical methods were used to pre-determine sample size. One pair of females was excluded from analysis due to a vaginal cytology sample that contained too few cells to accurately assess estrous state. All statistical analyses were carried out using R version 4.4.1 (R Core Team, 2024) and R Studio 2026.08.1+195 (Posit team, 2026).

### Data availability

All source data generated in this study will be deposited in a digital data repository, and this section will be modified prior to publication to include the persistent DOI for the dataset.

## Results

To test whether same-sex mounting by single-housed C57BL/6J female mice depends on estrous state, we conducted 20-minute-long social interaction trials between independent pairs of naturally cycling 3-days single-housed resident females and group-housed stimulus females (Fig. 1; n = 34 trials, n = 68 females). We refer to single-housed females as “residents”, because social interaction trials were conducted in that female’s home cage. Vaginal cytology was collected from both females on the day before and the day of the social interaction trial, and each female was categorized in a binary fashion as either receptive (in proestrus or estrus) or non-receptive (in metestrus or diestrus, see Methods) on the day of the social interaction trial. Overall, we found that single-housed resident females mounted group-housed stimulus females in 16 of 34 trials (∼47%), consistent with past work showing that short-term social isolation robustly promotes same-sex mounting in female mice (Zhao et al., 2021, 2025). We then tested whether the occurrence of same-sex mounting was coupled to the estrous state of either the single-housed resident female or the group-housed stimulus female.

**Figure 1.**
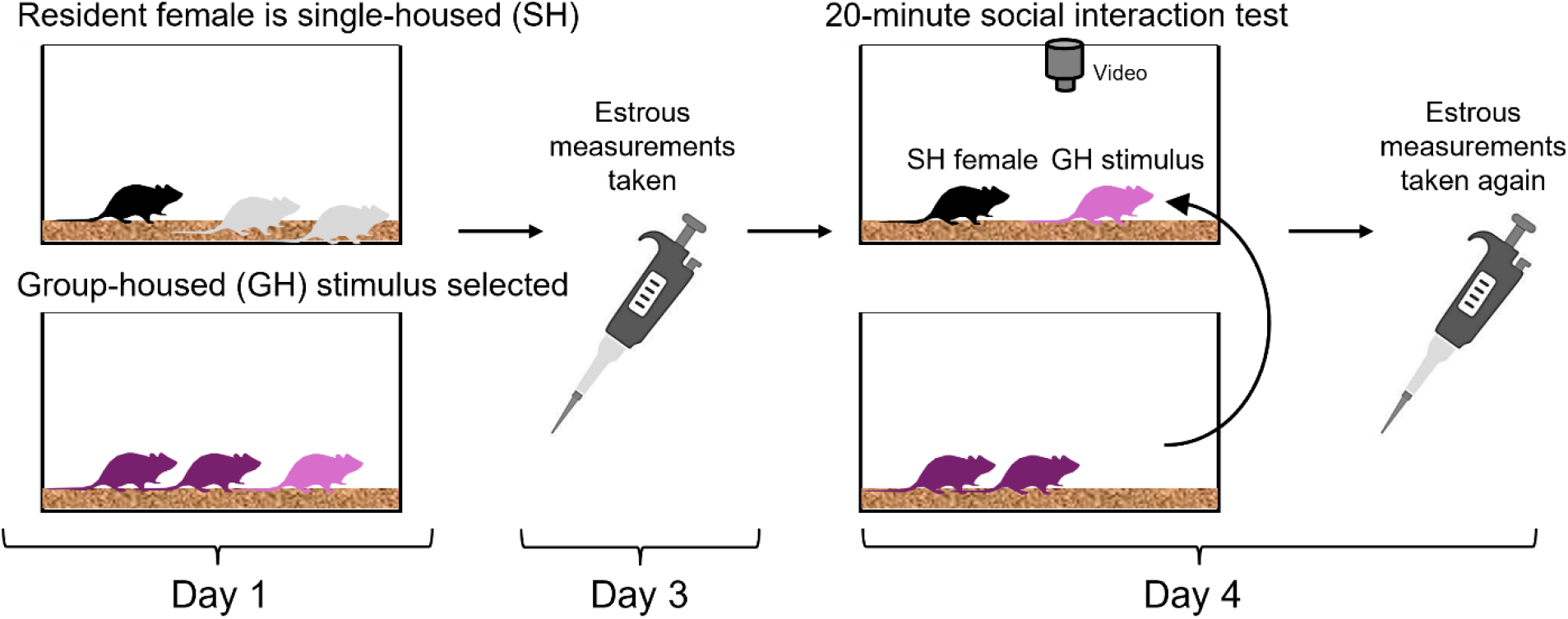
Experimental design. Schematic shows the timing of housing manipulations, estrous state measurements, and social interaction tests. SH, single-housed; GH, group-housed.

### Likelihood of mounting does not differ according to the estrous state of the single-housed resident female

We first considered whether displays of same-sex mounting depend on the estrous state of the single-housed resident female. Overall, 15 of 34 single-housed resident females were receptive on the day of behavioral testing (Fig. 2A). Given that we observed mounting in a similar proportion of the social interaction trials (16 of 34), one possibility is that same-sex mounting is displayed predominantly by receptive single-housed females. However, the proportion of total trials with mounting did not differ significantly according to the estrous state of the single-housed resident female (Fig. 2B; Fisher’s exact test, p = 1.00). This result shows that displays of same-sex mounting are not coupled to the estrous state of the single-housed resident female.

**Figure 2.**
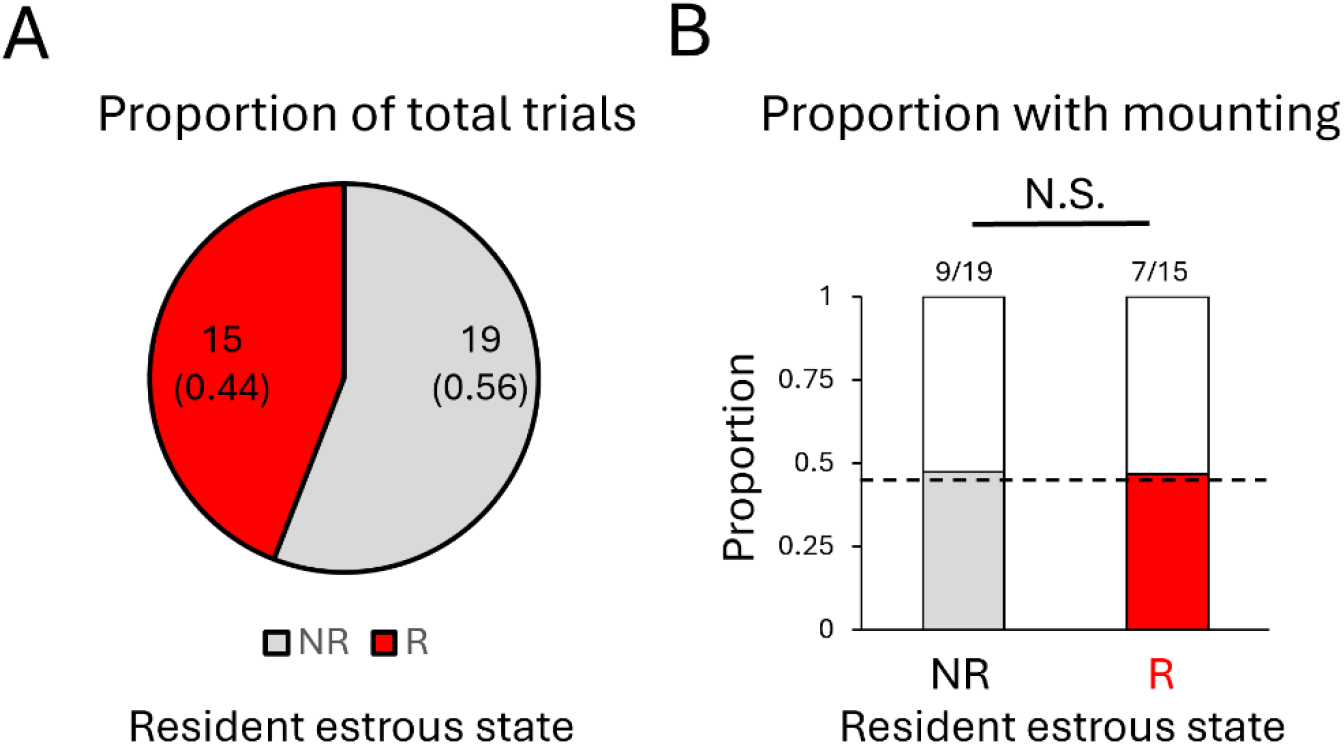
Likelihood of same-sex mounting according to estrous state of the single-housed resident female. (A) The total counts and proportion of trials with non-receptive (gray) and receptive (red) single-housed resident females are shown. (B) The proportion of trials with same-sex mounting is shown for trials that included a non-receptive (gray) or a receptive (red) single-housed resident female. The dotted line in (B) represents the overall proportion of trials with mounting in the dataset. NR, non-receptive; R, receptive.

### Single-housed resident females mount both receptive and non-receptive stimulus females

We next considered whether same-sex mounting depends on the estrous state of the group-housed stimulus female. Overall, relatively few trials included a group-housed stimulus female that was receptive on the day of behavioral testing (9 of 34 trials; Fig. 3A). We found that the proportion of trials with mounting did not differ significantly according to the estrous state of the GH stimulus female (Fig. 3B; Fisher’s exact test, p = 0.13), although we interpret this negative result cautiously due to the relatively small number of trials that included a receptive stimulus female in our dataset of naturally cycling female pairs. Nonetheless, we can conclude that same-sex mounting is observed in trials that included a receptive stimulus female as well as in trials that included a non-receptive stimulus female. These findings support the conclusion that displays of same-sex mounting by single-housed C57BL/6J females are not coupled to the estrous state of the stimulus female.

**Figure 3.**
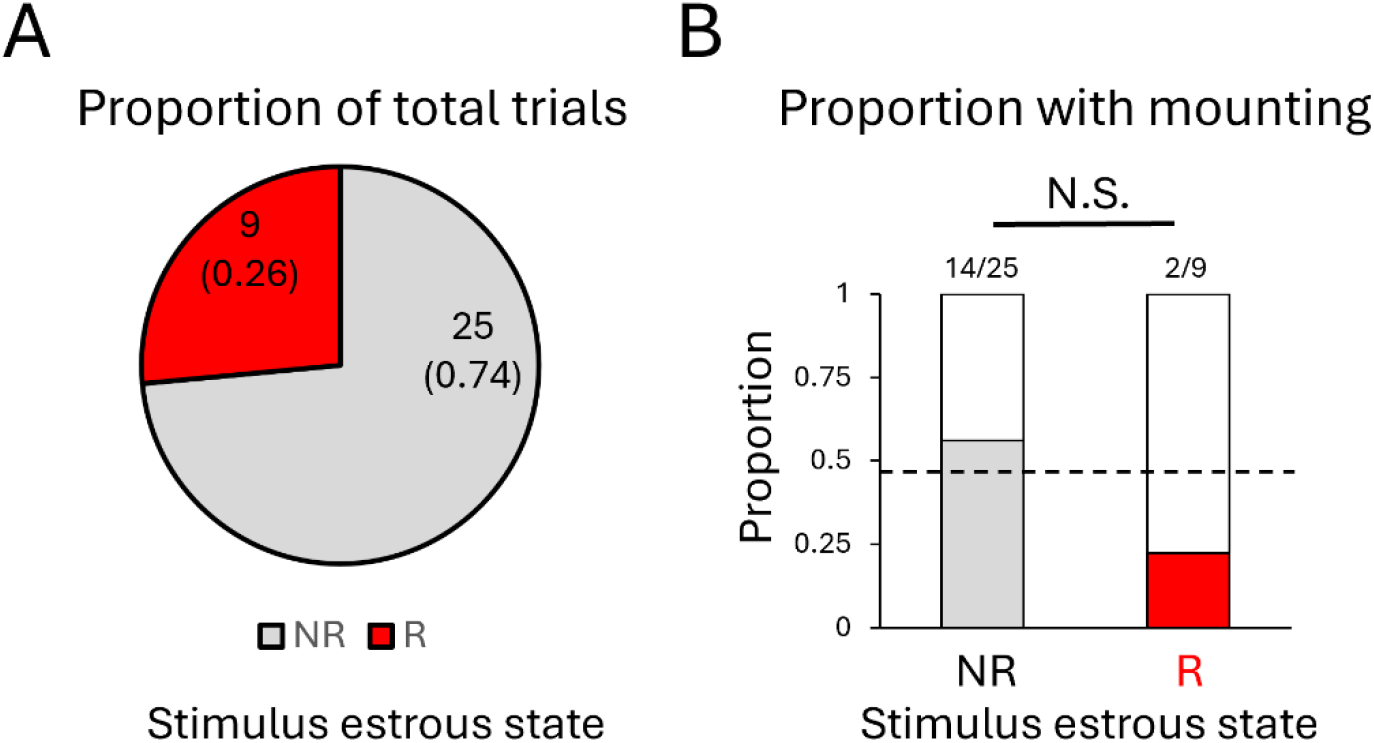
Likelihood of same-sex mounting according to estrous state of the group-housed stimulus female. (A) The total counts and proportion of trials with non-receptive (gray) and receptive (red) group-housed stimulus females are shown. (B) The proportion of trials with same-sex mounting is shown for trials that included a non-receptive (gray) or a receptive (red) group-housed stimulus female. The dotted line in (B) represents the overall proportion of trials with mounting in the dataset. NR, non-receptive; R, receptive.

### Same-sex mounting is observed in all resident-stimulus estrous state combinations

Given that same-sex mounting was observed in trials with both receptive and non-receptive single-housed resident females, as well as in trials with both receptive and non-receptive group-housed stimulus females, we finally asked whether same-sex mounting is observed in all resident-stimulus estrous state combinations. We first calculated the proportions of different resident-stimulus estrous state combinations across the trials in our dataset. On the day of behavioral testing, both females were non-receptive in about half of the social interaction trials (16 of 34, ∼47%), a non-receptive single-housed resident interacted with a receptive group-housed stimulus female in 3 of 34 trials (∼9%), a receptive single-housed resident interacted with a non-receptive group-housed stimulus female in 9 of 34 trials (∼26%), and both females were receptive in 6 of 34 trials (∼18%) (Fig. 4A).

**Figure 4.**
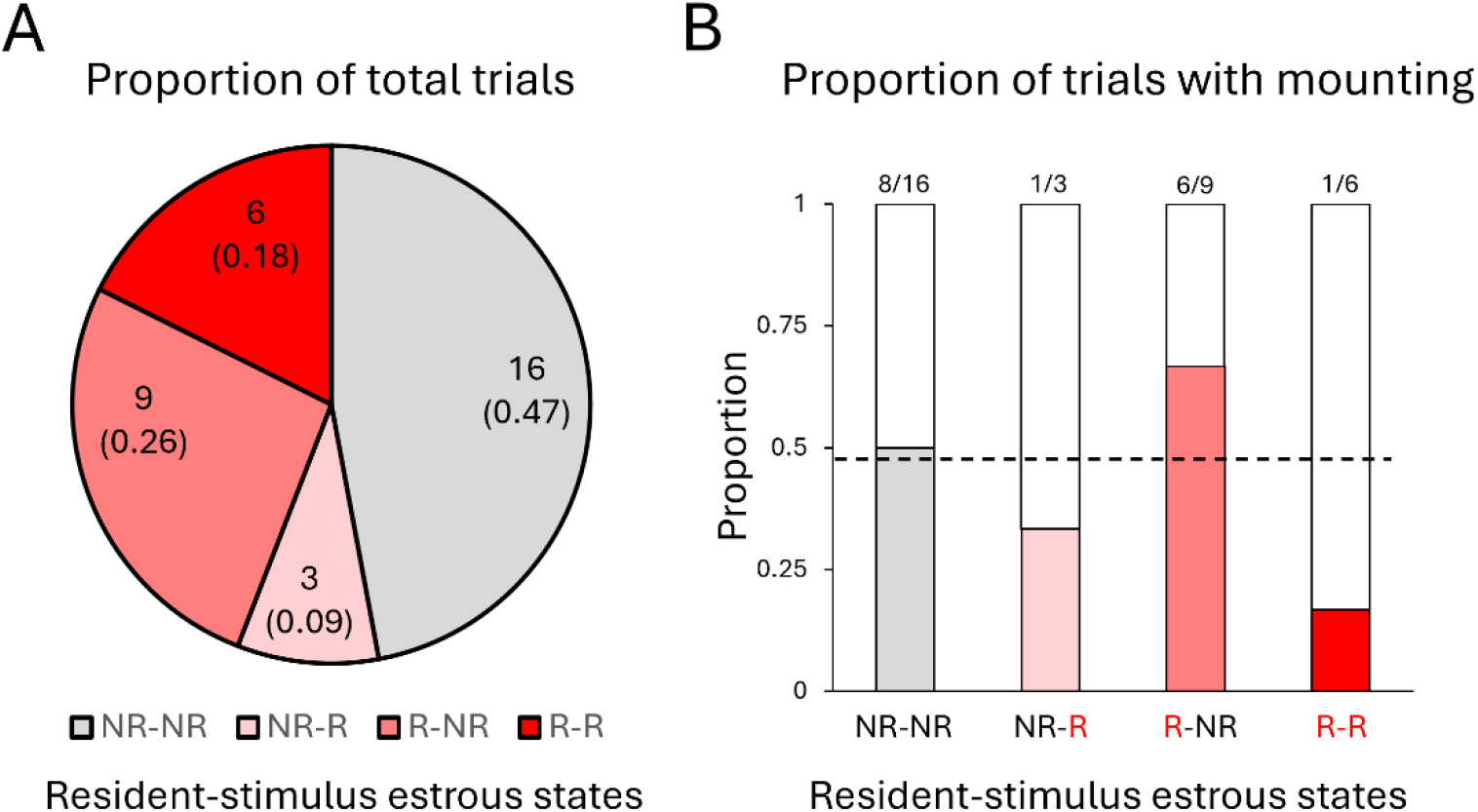
Likelihood of same-sex mounting according to resident-stimulus estrous state combinations. (A) The total numbers of social interaction trials with different resident-stimulus estrous state combinations are indicated. Parenthetical numbers indicate the proportion of total trials with each estrous state combination. (B) The proportion of trials with same-sex mounting is shown according to resident-stimulus estrous state combinations. The dotted line in (B) represents the overall proportion of trials with mounting in the dataset. NR, non-receptive; R, receptive.

The low probability of certain resident-stimulus estrous state combinations in naturally cycling female pairs precludes a formal comparison of the likelihood of mounting across all four resident-stimulus estrous state combinations. Nonetheless, we note that same-sex mounting was observed in at least one trial from each resident-stimulus estrous state combination (Fig. 4B). These data indicate that same-sex mounting by single-housed C57BL/6J females is not restricted to certain resident-stimulus estrous state combinations.

## Discussion

In the current study, we tested the relationship between female estrous state and same-sex mounting in single-housed C57BL/6J female mice during interactions with socially novel, group-housed stimulus females. We observed mounting in trials that included receptive and non-receptive single-housed residents, in trials that included receptive and non-receptive stimulus females, and even in trials from all resident-stimulus estrous state combinations. These findings do not support the conclusion that same-sex mounting in single-housed C57BL/6J female is coupled to estrous state.

We found that in naturally cycling female pairs, single-housed females were receptive in about half of the trials, and so our analyses included a large number of trials with receptive single-housed residents (n = 15) as well as a large number of trials with non-receptive single-housed residents (n = 19). The likelihood of mounting between these trial types was not different (Fig. 2B), indicating that estrous state of the single-housed female is not an important factor in determining the likelihood that the female displays same-sex mounting in this behavioral context.

Past work in rats found that sexually receptive (hormonally-primed, OVX) stimulus females were more likely to receive mounts that non-receptive stimulus females (Coniglio and Clemens, 1972; Fang and Clemens, 1999; Van de Poll et al., 1988). In partial agreement with these works, a recent study in large groups of outbred CD-1 female mice found that rates of same-sex mounting did not vary according to the estrous state of the female producing the mounts, but that females received higher rates of mounts when they were in estrus or metestrus as compared to diestrus (Williamson et al., 2019). In contrast, we found no significant difference in the likelihood of receiving mounts for receptive vs. non-receptive stimulus females, and if anything, we observed a tendency toward a reduced likelihood of mounting in trials that included receptive stimulus females (Figs. 3B, 4B). Given the relatively low chances of obtaining trials with receptive stimulus females using naturally cycling pairs (n = 9 of 34 trials), more work is needed to definitively test whether the likelihood of mounting differs according to stimulus female estrous state. Nonetheless, the fact that mounting was observed in trials with receptive stimulus females and in trials with non-receptive stimulus females supports the conclusion that same-sex mounting is not coupled to the estrous state of the stimulus female in this behavioral context.

If not estrous state, what factors play a major role in regulating the display of same-sex mounting in female rodents? Some studies have considered the possibility that female-female mounting serves to establish or maintain social hierarchies. Past work in pairs of female rats found that dominant females were more likely to mount than subordinate females, and that females were also more likely to mount socially novel females than familiar females (Fang and Clemens, 1999). More recent work in outbred CD-1 female mice housed in large social groups suggests that females use mounting alongside fighting and chasing to regulate dominance relationships (Williamson et al., 2019). Notably, the authors found that only a subset of females displayed mounting, but that when it occurred, mounting occurred 87% of the time in the direction of a more dominant female mounting a more subordinate female. Another interesting finding from this study is that although fighting occurred at the highest rates during the first 2 days of the experiment (i.e., during the initial formation of social hierarchy), mounting and chasing continued at similar rates throughout the two-week duration of the experiment. This finding suggests that outbred female mice might use mounting (and chasing) both to establish, as well as to maintain, social rank. It should be noted that both female rats and outbred female mice exhibit higher levels of aggression than the inbred female laboratory mice used in the current study. Nonetheless, more work is needed to test whether female-female mounting in single-housed C57BL/6J female mice correlates with social rank (either within the female’s original home cage or within newly formed social groups), is preferentially exhibited by only certain single-housed females, and/or varies according to the familiarity of a female social partner.

Although displays of same-sex mounting in single-housed C57BL/6J female mice are not coupled to female estrous state, it is still possible that hormonal signaling helps to mediate changes in female social behavior, including increasing the likelihood of same-sex mounting, following short-term social isolation. In addition to promoting the occurrence of same-sex mounting, three-days of single-housing also increases rates of social investigation and USV production in female mice (Zhao et al., 2021, 2025). Recent work identified a population of neurons in the preoptic hypothalamus (POA) that regulates this triad of changes in female social behavior following short-term social isolation (Zhao et al., 2025). Moreover, past work has implicated the POA, and Esr1-expressing POA neurons more specifically, in regulating the production of male-female mounting during rodent courtship interactions (Floody, 1989; Karigo et al., 2021; Wei et al., 2018). Together, these studies raise the possibility that hormonal signaling within the POA might promote the display of female-female mounting following social isolation, although this idea remains to be explored in future studies.

## Acknowledgements

We thank the CARE staff for their excellent mouse husbandry. This research did not receive any specific grant from funding agencies in the public, commercial, or not-for-profit sectors.

## References

Bailey, N.W., Zuk, M., 2009. Same-sex sexual behavior and evolution. Trends Ecol. Evol. 24, 439–446.

Bakker, J., De Mees, C., Douhard, Q., Balthazart, J., Gabant, P., Szpirer, J., Szpirer, C., 2006. Alpha-fetoprotein protects the developing female mouse brain from masculinization and defeminization by estrogens. Nat. Neurosci. 9, 220–226.

Bakker, J., Honda, S.-I., Harada, N., Balthazart, J., 2002. The Aromatase Knock-Out Mouse Provides New Evidence That Estradiol Is Required during Development in the Female for the Expression of Sociosexual Behaviors in Adulthood. J. Neurosci. 22, 9104–9112.

Baum, M.J., Södersten, P., Vreeburg, J.T.M., 1974. Mounting and receptive behavior in the ovariectomized female rat: Influence of estradiol, dihydrotestosterone, and genital anesthetization. Horm. Behav. 5, 175–190.

Beach, F.A., Rasquin, P, 1942. Masculine copulatory behavior in intact and castrated female rats. Endocrinology. 31, 393–409.

Beach, F. A., 1968. Factors involved in the control of mounting behavior by female mammals, in Diamond, M. (Ed.), Perspectives of Reproduction and Sexual Behavior: A Memorial to William C. Young. Indiana University Press, Bloomington, pp. 83–131.

Coniglio, L., Clemens, L.G., 1972. Stimulus and experiential factors controlling mounting behavior in the female rat. Physiol. Behav. 9, 263–267.

Dagg, A.I., 1984. Homosexual behaviour and female-male mounting in mammals—a first survey. Mamm. Rev. 14, 155–185.

de Jonge, F.H., Burger, J., Van de Poll, N.E., 1986. Variable mounting levels in the female rat: The influence of experience and acute effects of testosterone. Behav. Brain Res. 20, 39–46.

de Jonge, F.H., Burger, J., Van Haaren, F., Overdijk, H., Van De Poll, N.E., 1987. Sexual experience and preference for males or females in the female rat. Behav. Neural Biol. 47, 369–383.

Edwards, D.A., Burge, K.G., 1971. Estrogenic arousal of aggressive behavior and masculine sexual behavior in male and female mice. Horm. Behav. 2, 239–245.

Fang, J., Clemens, L.G., 1999. Contextual determinants of female-female mounting in laboratory rats. Anim. Behav. 57, 545–555.

Floody, O.R., 1989. Dissociation of hypothalamic effects on ultrasound production and copulation. Physiol. Behav. 46, 299–307.

Gómez, J.M., Gónzalez-Megías, A., Verdú, M., 2023. The evolution of same-sex sexual behaviour in mammals. Nat. Commun. 14, 5719.

Hashikawa, K., Hashikawa, Y., Tremblay, R., Zhang, J., Feng, J.E., Sabol, A., Piper, W.T., Lee, H., Rudy, B., Lin, D., 2017. Esr1+ cells in the ventromedial hypothalamus control female aggression. Nat. Neurosci. 20, 1580–1590.

Karigo, T., Kennedy, A., Yang, B., Liu, M., Tai, D., Wahle, I.A., Anderson, D.J., 2021. Distinct hypothalamic control of same- and opposite-sex mounting behaviour in mice. Nature 589, 258–263.

Martel, K.L., Baum, M.J., 2009. Adult Testosterone Treatment But Not Surgical Disruption of Vomeronasal Function Augments Male-Typical Sexual Behavior in Female Mice. J. Neurosci. 29, 7658–7666.

Noble, R., 1974. Estrogen plus androgen induced mounting in adult female hamsters. Horm. Behav. 5, 227–234.

Noble, R.G., 1977. Mounting in female hamsters: Effects of different hormone regimens. Physiol. Behav. 19, 519–526.

Posit team. (2026). RStudio: Integrated Development Environment for R. Posit Software, PBC. http://www.posit.co/

R Core Team. (2024). R: A Language and Environment for Statistical Computing. R Foundation for Statistical Computing. https://www.R-project.org/

Södersten, P., 1972. Mounting behavior in the female rat during the estrous cycle, after ovariectomy, and after estrogen or testosterone administration. Horm. Behav. 3, 307–320.

Van de Poll, N.E., Taminiau, M.S., Endert, E., Louwerse, A.L., 1988. Gonadal steroid influence upon sexual and aggressive behavior of female rats. Int. J. Neurosci. 41, 271–286.

Wei, Y.-C., Wang, S.-R., Jiao, Z.-L., Zhang, W., Lin, J.-K., Li, X.-Y., Li, S.-S., Zhang, X., Xu, X.-H., 2018. Medial preoptic area in mice is capable of mediating sexually dimorphic behaviors regardless of gender. Nat. Commun. 9, 279.

Wersinger, S.R., Sannen, K., Villalba, C., Lubahn, D.B., Rissman, E.F., De Vries, G.J., 1997. Masculine Sexual Behavior Is Disrupted in Male and Female Mice Lacking a Functional Estrogen Receptor α Gene. Horm. Behav. 32, 176–183.

Williamson, C.M., Lee, W., DeCasien, A.R., Lanham, A., Romeo, R.D., Curley, J.P., 2019. Social hierarchy position in female mice is associated with plasma corticosterone levels and hypothalamic gene expression. Sci. Rep. 9, 7324.

Zhao, X., Chae, Y., Smith, D., Chen, V., DeFelipe, D., Sokol, J.W., Sadangi, A., Tschida, K., 2025. Short-term social isolation acts on hypothalamic neurons to promote social behavior in a sex- and context-dependent manner. eLife 13, RP94924.

Zhao, X., Ziobro, P., Pranic, N.M., Chu, S., Rabinovich, S., Chan, W., Zhao, J., Kornbrek, C., He, Z., Tschida, K.A., 2021. Sex- and context-dependent effects of acute isolation on vocal and non-vocal social behaviors in mice. PLoS One 16, e0255640.

